# A Dimensional Approach to Assessing Psychiatric Risk in Adults Born Very Preterm

**DOI:** 10.1101/140772

**Authors:** Jasmin Kroll, Sean Froudist-Walsh, Philip J. Brittain, Chieh-En Jane Tseng, Vyacheslav Karolis, Robin M Murray, Chiara Nosarti

**Author notes:** Address correspondence to: Jasmin Kroll; Department of Psychosis Studies, Institute of Psychiatry, Psychology and Neuroscience, King’s College London, 16 De Crespigny Park, London SE5 8AF, UK.

## Abstract

**Background:** Individuals who were born very preterm have higher rates of psychiatric diagnoses compared to term-born controls; however, it remains unclear whether they also display increased sub-clinical psychiatric symptomatology. Hence our objective is to utilise a dimensional approach to assess psychiatric symptomatology in adults who were born very preterm.

**Methods:** 152 adults who were born very preterm (before 33 weeks’ gestation; gestational range 24–32 weeks) and 96 term-born controls. We examined participants’ clinical profile using the Comprehensive Assessment of At-Risk Mental States (CAARMS), a measure of sub-clinical symptomatology that yields seven subscales including general psychopathology, positive, negative, cognitive, behavioural, motor and emotional symptoms, in addition to a total psychopathology score. Intellectual abilities were examined using the Wechsler Abbreviated Scale of Intelligence.

**Results:** Between-group differences on the CAARMS showed elevated symptomatology in very preterm participants compared to controls in positive, negative, cognitive and behavioural symptoms. Total psychopathology scores were significantly correlated with IQ in the very preterm group only. In order to examine the characteristics of participants’ clinical profile a principal component analysis was conducted. This revealed two components, one reflecting a non-specific psychopathology dimension, and the other indicating a variance in symptomatology along a positive-to-negative symptom axis. K-means (k=4) were used to further separate the study sample into clusters. Very preterm adults were more likely to belong to the high non-specific psychopathology cluster compared to controls.

**Conclusion and Relevance:** Very preterm individuals demonstrated elevated psychopathology compared to full-term controls. Psychiatric risk was characterised by a non-specific clinical profile and was associated with lower IQ.

## Introduction

Within a conceptual framework suggesting that mental illness lies on a continuum with typical behavioural traits, (Hack *et al.*, 2004b) psychiatric vulnerability can be measured in both healthy and clinical samples, (van Os *et al.*, 2009) as well as in ‘high-risk’ populations (Demjaha *et al.*, 2012). Approximately 25% of children born before 32 weeks (i.e., very preterm) have persisting neuropsychiatric concerns, which are characterized by inattention, anxiety, socio-emotional difficulties, and internalizing problems (Johnson and Marlow, 2011). Prevalence rates of emotional and behavioural problems range between 8% and 39%, depending on gestational age, with the most immature children being at greatest risk of impairment (Arpi and Ferrari, 2013). This ‘behavioural phenotype’ results in increased rates of sub-threshold symptomatology that, at the furthest end of the distribution, become clinically significant (Hack *et al.*, 2009, Johnson and Marlow, 2011). Very preterm children are thus at higher risk than controls of developing autism spectrum disorder, attention deficit hyperactivity disorder and anxiety disorder (Johnson and Marlow, 2011, Treyvaud *et al.*, 2013b). In adult life, preterm born individuals report elevated levels of psychological distress, (Wiles *et al.*, 2005) continue to be vulnerable to mental health problems (Hack *et al.*, 2004a) and are between 2.5 and 7.4 times at greater risk of being hospitalized with a range of psychiatric disorders compared to controls, as indicated in population-based studies (Johnson and Marlow, 2011, Nosarti *et al.*, 2012a, Nosarti *et al.*, 2012c, Treyvaud *et al.*, 2013a). However, it is likely that the clinical presentation of very preterm individuals will extend across the standard diagnostic boundaries (Johnson and Marlow, 2014), and therefore be overlooked in prevalence studies. Indeed, individuals who were born very preterm often show qualitative differences in their psychiatric presentation, (Johnson and Marlow, 2011, Johnson and Wolke, 2013) suggesting that they may have a different aetiological risk and symptom profile from that seen in the general population.

The lack of specificity in outcome suggests that preterm birth may represent a risk factor for various types of psychopathology, possibly due to developmental alterations in whole brain connectivity preferentially affecting corticostriatal and thalamocortical connections, (Ball *et al.*, 2015, Fischi-Gomez *et al.*, 2014) which have also been described in psychiatric conditions with a neurodevelopmental component (e.g. (Arpi and Ferrari, 2013, Ball *et al.*, 2015, Fischi-Gomez *et al.*, 2014, Johnson and Marlow, 2011)). Furthermore, structural brain alterations are observed in individuals who are at high risk of developing a psychiatric disorder and even in those individuals *who do not* subsequently receive a clinical diagnosis (Takahashi *et al.*, 2009). The study of vulnerable individuals with elevated psychiatric symptomatology without a clinical diagnosis represents an unrecognized public health concern as they ‘suffer in silence’ reporting a lower quality of life (Carta and Angst, 2016) and distress-impairment (McLaughlin *et al.*, 2015).

Based on existing evidence suggesting that very preterm individuals have an increased risk of both sub-threshold psychiatric symptomatology and clinical disorders, the aim of this study was to utilise a dimensional approach to examine whether adults who were born very preterm would demonstrate elevated levels of psychopathology compared to controls, and secondly, to explore the specific characteristics of their symptom profile.

## Materials and Methods

### Study population

152 individuals who were born before 33 weeks’ gestation between 1979 and 1984 and were admitted to the neonatal unit of University College Hospital (UCH) London within five days of birth were recruited for this study. Participants were then entered into a follow-up study and were reassessed periodically throughout their lives (Froudist-Walsh *et al.*, 2015, Nam *et al.*, 2015, Stewart *et al.*, 1989). Neonatal variables collected at birth included: birth weight, gestational age and severity of perinatal brain injury, based on neonatal cranial ultrasound classification summarized as a) normal, no-periventricular haemorrhage (no-PVH), b) uncomplicated periventricular haemorrhage without ventricular dilatation (PVH), and c) periventricular haemorrhage with ventricular dilatation (PVH+DIL) (Nosarti *et al.*, 2011b) (Nosarti *et al.*, 2011a).

Very preterm individuals who were assessed at the current follow-up did not differ significantly from those who were not assessed in terms of their birth weight (Assessed at 30: 1305.83 grams, Not assessed at 30: 1371.75 grams, t=−1.78, df=450, p=.075), however, those who were assessed were born at a slightly younger gestational age than those who were not (Assessed at 30: mean gestational age = 29.18 weeks, Not assessed at 30: mean gestational age=29.67, t=−2.23, df=451, p=.026) and there was a higher proportion of males in the returning cohort (Assessed at 30: 62% male, Not assessed at 30: 48% male, X^2^=7.19, df=1, p=<0.01).

The term-born control group consisted of 96 individuals recruited from advertisements in the local community. Inclusion criteria were full-term birth (38–42 weeks) and birth weight >2500 grams. Exclusion criteria were a history of neurological conditions including meningitis, head injury and cerebral infections.

The study was undertaken with the understanding and written consent of each subject, with the approval of the appropriate local ethics committee, and in compliance with national legislation and the Code of Ethical Principles for Medical Research Involving Human Subjects of the World Medical Association (Declaration of Helsinki).

### Socio-demographic, cognitive and behavioural assessment

Participants’ socio-economic status (SES) was assessed with Her Majesty’s Stationary Office Standard Occupational Classification Information (Her Majesty’s Stationary Office, 1991). IQ was examined using the Wechsler Abbreviated Scale of Intelligence (WASI; (Wechsler, 1999).

Psychiatric symptomatology was assessed with the ‘Comprehensive Assessment of At-Risk Mental States’ (CAARMS; (Yung *et al.*, 2005). The CAARMS is an interviewer-rated, semistructured tool, measuring current rates of psychopathology on the following subscales: positive and negative symptoms, cognitive problems, emotional disturbance, behavioural changes, motor/physical changes and general psychopathology. General psychopathology included depression, anxiety, mania, and mood swings. Each scale is rated on a 0–6 severity scale (‘0 – Never/absent’ to ‘6 – Extreme’). Inter-rater reliability was assessed by comparing ratings for all subscales for three very preterm individuals who were assessed by both study raters and a ‘gold-standard’ rater (an experienced psychiatrist). Intra-class Correlation Coefficients were 0.89 between study raters PJB and JK and .90 and .86 between raters PJB and JK and the gold-standard rater respectively. These values represent ‘Almost Perfect’ agreement (Landis and Koch, 1977).

### Statistical analysis

Matlab, version R2016a (Mathworks, MA, USA) and SPSS for Macintosh, version 22.0 (IBM, Armonk, NY), were used for the statistical analyses. Group differences in sociodemographic measures were examined using independent *t*-test or Chi-Square test, with significance set at p<0.05.

#### Part 1: Group differences in symptomatology

Between-group differences on each of the CAARMS subscales were explored using the Mann-Whitney U-test. Spearman correlation was used to examine the association between IQ and Total Psychopathology.

To further examine between-group differences, a previously described cut-off was used to define individuals born very preterm that are at risk of clinically significant problems, defined as a CAARMS score greater than or equal to the 90^th^ percentile score of controls (Healy *et al.*, 2013) (Rickards *et al.*, 2001). This group will be referred to as ‘high-risk’ in the text.

In order to quantify this risk on each CAARMS scale, Fisher’s Exact Test was performed and summarized as Odds Ratio. Motor symptoms were excluded from the analyses as very few controls scored above zero on this measure. Multiple comparison correction was performed using false discovery rate (FDR) correction (Benjamini and Hochberg, 1995).

#### Part 2: Specificity of symptom profile

A principal component analysis (PCA) was performed on the CAARMS scales to provide a dimensional overview of the pattern of psychiatric symptoms. A skree plot was used to identify the number of components that parsimoniously described the variance in the psychopathology data. Once the ideal number of principal components was found, k-means clustering was performed, in order to group individuals according to their symptom distribution. In order to analyse whether very preterm born individuals were more likely to experience a certain cluster of symptoms compared to controls, a Chi-square test was used. Pairwise Fisher’s Exact Test was performed to compare participants’ symptoms distribution in each cluster to that in the low psychopathology cluster.

## Results

**Neonatal, socio-demographic, cognitive** variables **and psychiatric history** are presented in Table 1. There were more men than women in the very preterm group compared to controls. Very preterm individuals had a significantly lower IQ and were more likely to report a lifetime psychiatric history compared to controls.

**Table 1:** Neonatal, socio-demographic, cognitive variables and psychiatric history for term-born and very preterm participants.

|  | Term (n=96) | Very preterm (n=152) | Statistics |
| --- | --- | --- | --- |
| Gestational age (weeks) | - | 29.28 (SD 2.09) | - |
| Birth weight (grams) | - | 1312.45 (SD 349.92) | - |
| Neonatal Cranial Ultrasound Classification<br>(% no-PVH/PVH/PVH+DIL) | - | 49/22/28 | - |
| Age (years) | 30.64 (5.24) | 31.46 (2.33) | N.S. |
| Gender (% male) | 46 | 59 | $\chi^2_{(1)} = 4.24; p = .04$ |
| Full-scale IQ | 111.62 (13.15) | 102.40(15.27) | t = 4.42 p = .000 |
| Self-reported psychiatric history | 10 (11.4) | 40 (26.5) | $\chi^2_{(1)} = 7.69$; p=.006 |

no-PVH: normal neonatal cranial ultrasound, PVH: uncomplicated periventricular haemorrhage without ventricular dilatation, PVH+DIL: periventricular haemorrhage with ventricular dilatation. Means and standard deviations (SD) are presented, unless specified.

### CAARMS results

Very preterm participants had significantly elevated levels of emotional disturbances, positive, negative, cognitive, negative and motor symptoms compared to controls (Table 2), and all results survived FDR correction. However, there were no significant between-group differences in emotional disturbance, while differences in general psychopathology reached borderline levels of significance.

**Table 2:** CAARMS scores for very preterm and term-born participants.

| **CAARMS measures**<br>**Mean** | **Term** | **VPT** | **Mann-Whitney U** | **p** |
| --- | --- | --- | --- | --- |
| Total Psychopathology | 7.57 | 15.39 | 4179 | **.001** |
| Positive Symptoms | .056 | 1.74 | 4448 | **.002** |
| Cognitive Symptoms | 1.02 | 2.04 | 4376 | **.003** |
| Emotional Disturbances | 0.77 | 1.21 | 5265 | .414 |
| Negative Symptoms | 0.93 | 1.96 | 4717.5 | **.020** |
| Behavioural Changes | 1.21 | 2.56 | 4825 | **.042** |
| General Psychopathology | 2.89 | 4.89 | 4799.5 | .051 |

CAARMS: Comprehensive Assessment of At-risk Mental States. Total psychopathology score is the sum of all subscales.

In the whole sample, higher Total Psychopathology scores were significantly associated with lower full-scale IQ (Spearman’s r= −.268, p=.000); however within group analyses showed that this association was significant in the very preterm group (r= −.259; p=.003), but not in controls (r= −.187; p=.114). The difference between these two correlation coefficients was not statistically significant (Fisher’s z=−0.51, p>0.05).

### Group and ‘high-risk’ symptomatology

Very preterm individuals were significantly more likely than controls to score above the ‘high-risk’ threshold for total symptomatology (p= 0.044), as well as for the positive (p = 0.002), cognitive (p =0.002) and negative (p =0.014) scales. There were no significant differences between the groups on the emotional disturbance (p =0.596), behavioural (p =0.057), or general psychopathology (p =0.101) scales as described in Figure 1. After FDR correction significant between group results remained on the positive, cognitive and negative scales.

**Figure 1:**
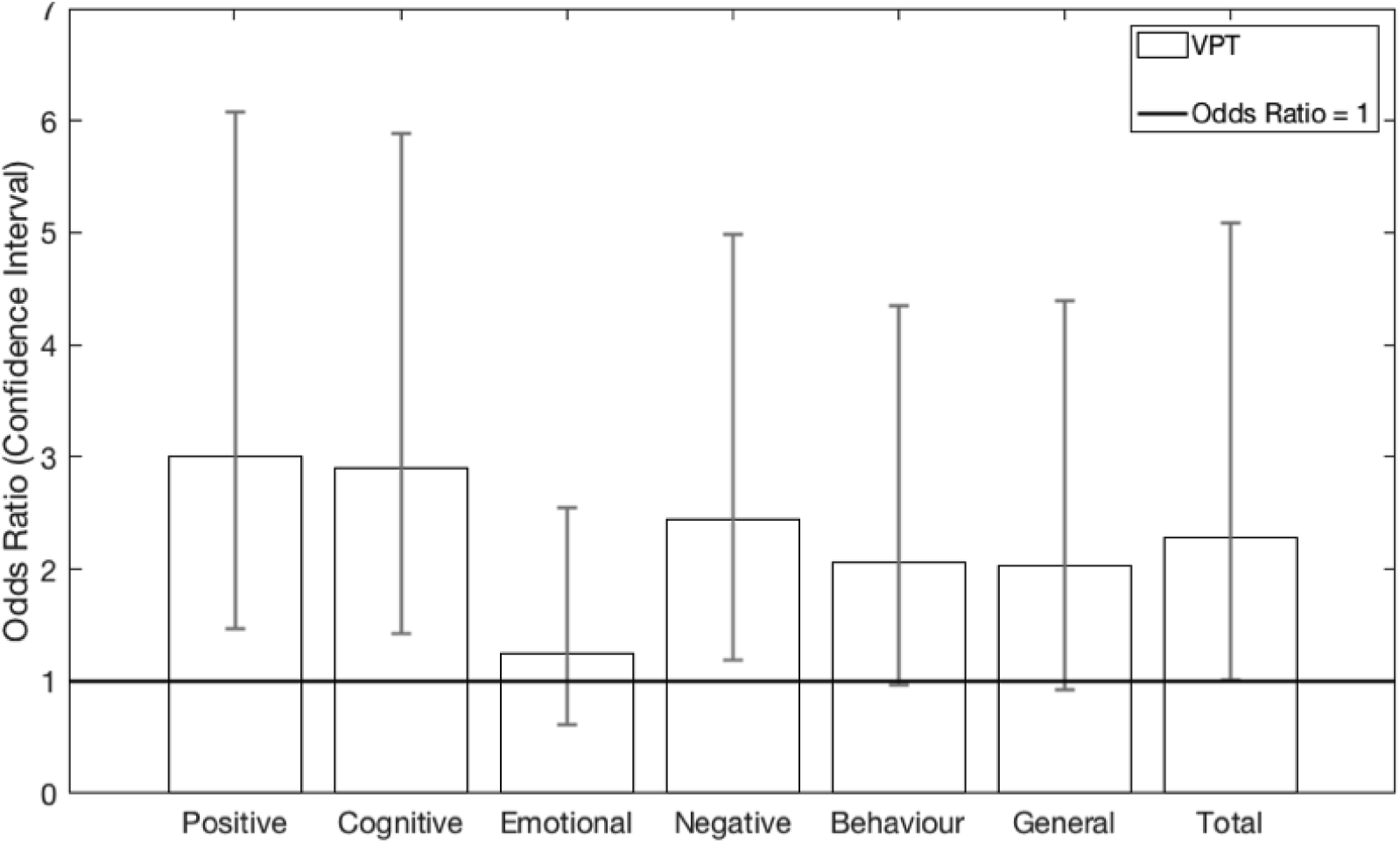
Odds ratio for being at ‘high-risk’ in adults born very preterm. Adults born very preterm were more likely than controls to belong to the ‘high-risk’ category, on the basis of total symptoms, as well as positive, negative and cognitive symptoms.

### Symptom clustering

Principal components analysis revealed two components that explained 77.44% of the variance in the CAARMS scales (principal component 1 (PC1) = 67.08%, principal component 2 (PC2) = 10.36%). PC1 had negative weights of a similar size (between −0.38 and −0.43) for each CAARMS subscale, indicating a non-specific psychopathology dimension. PC2 had large positive weightings on positive and cognitive subscales (0.57, 0.56) and relatively large negative weightings on the negative and behavioural subscales (−0.32, −0.45), indicating a variance in symptomatology along a positive-to-negative symptom axis.

In order to investigate if very preterm birth was likely to be a risk factor for a specific psychiatric dimension, we used K-means clusters (k=4) to separate the study sample into clusters that differed on their loadings on both the non-specific psychopathology axis, and the positive-to-negative symptom axis. Specifically, Cluster 1 contained individuals who scored high on non-specific psychopathology. Cluster 2 contained individuals who scored low on non-specific psychopathology. Clusters 2 and 3 both exhibited only mild overall symptoms, but were separated on the positive-to-negative axis, with individuals in Cluster 2 tending to have more positive and cognitive symptoms, and individuals in Cluster 3 tending to have more negative and behavioural symptoms (Figure 2).

**Figure 2:**
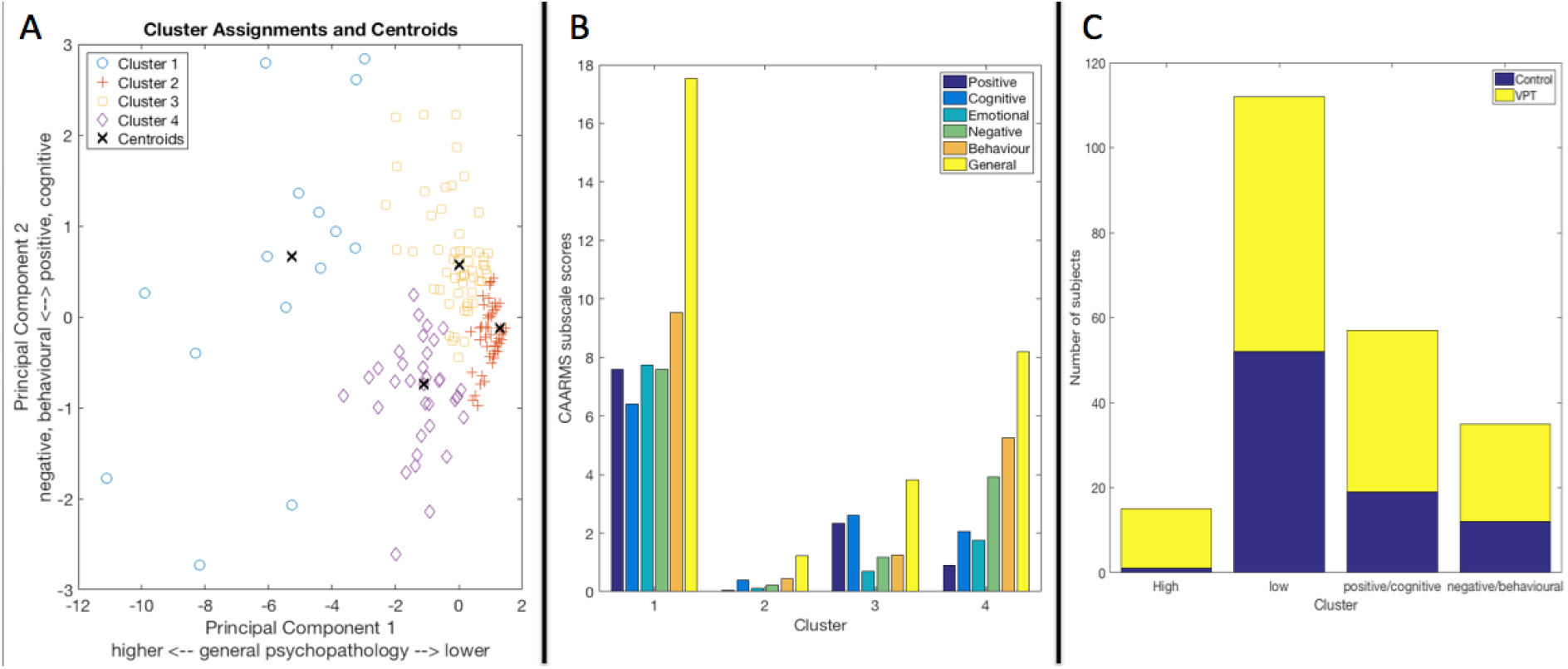
Symptom clustering. A) Principal Components Analysis revealed two major components. The first component separated individuals with high and low non-specific psychopathology. The second component accounted for variance along a positive/cognitive to negative/behaviour symptom axis. K-means clustering of participants’ loadings on these two components identified four psychopathology clusters. B) CAARMS sub-scores by clusters: Cluster 1: high non-specific psychopathology; Cluster 2: low non-specific psychopathology; Cluster 3: high positive and cognitive symptoms; Cluster 4: high negative and behavioural symptoms. C) Group composition by cluster.

The distribution of the groups within each cluster is shown in Table 3 and Figure 2C. A Chisquare test indicated significant between group differences in their distribution into clusters (X^2^ = 10.31, p = .016). In order to further probe whether study participants were more likely to belong to a particular psychopathology cluster than chance, we performed a series of Fisher’s Exact Tests to study whether the prevalence of very preterm participants in the high non-specific psychopathology, positive/cognitive and negative/behavioural clusters was greater than their prevalence in the low general psychopathology cluster. Results indicated that preterm individuals were more likely to belong to the high non-specific psychopathology cluster than controls, but this was not found for the positive/cognitive or the negative/behavioural cluster (Table 2).

**Table 3:** Distribution of controls and adults born very preterm within psychopathology clusters

| Cluster | Control<br>N (%) | VPT<br>N (%) | Odds Ratio<br>[95% Confidence Interval] | p |
| --- | --- | --- | --- | --- |
| 1. Low non-specific psychopathology | 52 (61.9%) | 60 (44.4%) | – | – |
| 1. High non-specific psychopathology | 1 (1.2%) | 14 (10.4%) | 12.13 [1.54, 95.43] | 0.0039 |
| 3. High positive/cognitive symptoms | 19 (22.6%) | 38 (28.2%) | 1.73 [0.89, 3.37] | 0.137 |
| 4. High negative/behavioural symptoms | 12 (14.3%) | 23 (17.0%) | 1.66 [0.75, 3.66] | 0.244 |

## Discussion

The current study found that adults who were born very preterm demonstrated elevated psychiatric symptomatology compared to controls. As well as displaying increased total psychopathology, they showed increased positive, cognitive and negative symptoms. Individuals who were born very preterm were also between one- to three-fold more likely than controls to belong to a ‘high-risk’ group (defined by CAARMS scores above the 90^th^ percentile of control scores) on several symptom scales.

These results are in line with previous research, indicating higher rates of psychiatric symptomatology in very preterm children, adolescents and in young adults’(Hack *et al.*, 2004a, Healy *et al.*, 2013, Johnson *et al.*, 2010). Although the instrument we used, the CAARMS, was designed to explore subclinical psychopathology believed to indicate an imminent development of first-episode psychosis, it covers wider psychopathological domains and in this respect our results could be comparable with population-linkage studies that reported a significant association between very preterm birth and a number of psychiatric disorders such as depression, anxiety, schizophrenia and bipolar affective disorder (Johnson and Marlow, 2011) (Nosarti *et al.*, 2012b) (Nosarti *et al.*, 2012a) (Crump *et al.*, 2010). Hence, the findings presented here suggest the existence of a major, yet poorly appreciated, psychiatric burden in adults who were born very preterm.

### Participants’ Clinical Profile

In the current assessment very preterm individuals scored higher on the majority of CAARMS sub-scales compared to controls, which may suggest a non-specific risk (Nosarti *et al.*, 2012b). Nonetheless, several of the symptoms that have been previously described as characterising a ‘preterm behavioural phenotype’ in childhood (Johnson and Marlow, 2011) and that are included in the CAARMS continued to be prevalent in our very preterm sample in adult life and these included attention and concentration difficulties, social withdrawal, cognitive changes, alogia, anhedonia, a decreased ability to perform adult roles, apathy, and depression/anxiety. In this sense, such symptom profile may transcend current diagnostic boundaries.

One challenge in understanding the psychiatric profile of adults who were born very preterm, is to disentangle the commonly described cognitive deficits, such as IQ and executive function deficits, which are thought to underlie social and behavioural problems (Aarnoudse-Moens *et al.*, 2009, Anderson and Doyle, 2004, Delobel-Ayoub *et al.*, 2009). Considering the significant association between IQ and psychiatric symptomatology, we are tempted to speculate that preterm adults may be represent an aetiologically and prognostically distinct subgroup characterised by cognitive impairments (Fusar-Poli *et al.*, 2012). Moreover, prospective studies indicate that in populations at risk of developing psychiatric disorders, deficits in social cognition and executive function, along with emotional and behavioural disturbances, may arise in childhood (van Os and Kapur, 2009) and continue into adulthood, when symptom expression may change in magnitude and character to reflect age-related changes (Hudziak *et al.*, 2007).

Indeed, a study conducted in a partially overlapping subsample of the current cohort in mid-adolescence, reported elevated scores on the ‘Social Problems’ scale of the parent-rated Child Behaviour Checklist (CBCL; (Healy *et al.*, 2013), with items such as “does not get along with peers”, “gets teased” and “too dependent”. Similarly, at age 18, this cohort was found to have increased levels of psychiatric ‘caseness’ according to the Clinical Interview Schedule – Revised (Walshe *et al.*, 2008) with the most common diagnoses being mood and anxiety disorders. It may be, therefore, that these results represent a continuum of psychiatric risk from mid-adolescence through to adulthood, albeit highlighted with different instruments.

### Neurodevelopmental Origin of Psychiatric Risk

The current findings support the notion of a neurodevelopmental origin of psychiatric disorder (Howes and Murray, 2014). However, the precise pathway linking very preterm birth and psychopathology remains unclear. We previously proposed a theoretical framework, which considered both biological and environmental contributions (Montagna and Nosarti, 2016). According to this model, very preterm birth leads to long-lasting structural and functional brain alterations in socio-emotional and cognitive networks (Fischi-Gomez *et al.*, 2014, Papini *et al.*, 2016, Reininghaus *et al.*, 2016). These may increase an individual’s vulnerability to psychopathology, including enhanced stress sensitivity and aberrant salience (Reininghaus *et al.*, 2016). Furthermore, very preterm individuals may be particularly susceptible to bullying, social defeat and internalising symptoms, which have also been studied as risk factors for psychopathology (Johnson and Marlow, 2011, Montagna and Nosarti, 2016, Valmaggia *et al.*, 2015, Wolke *et al.*, 2015). Within this theoretical framework, psychiatric disorder may represent the endpoint of a risk pathway that beings at birth (Dutta *et al.*, 2007). Hence our findings highlight the importance of collecting perinatal data as part of routine psychiatric assessments, of monitoring possible antecedents to psychiatric disorder in preterm born individuals and of developing preventative interventions early in life. Moreover, further studies are required to examine the generalizability of the current results to other high-risk populations, such as those with obstetric complications other than very preterm birth and those at genetic risk for psychopathology.

### Limitations

The current study had a number of limitations. Our study participants were born in the late 1970’s and early 1980’s and, due to advances in neonatal care, may have displayed mental health symptoms in adulthood, which are not representative of very preterm cohorts born in more recent years. Similar to other longitudinal studies, attrition is a critical limitation; participants studied here were a subset of the original cohort. A previous study found a bias in selective dropout where those with the worst outcomes did not return for assessments; however, this would decrease the prevalence of self-reported psychiatric history, indicating our results may be an underestimation of participants’ current psychiatric profile (Wolke *et al.*, 2009).

We further acknowledge that a major limitation of this study is the use of one assessment tool, which was originally designed to evaluate attenuated symptomatology in individuals at risk of psychosis. Considering the overlap between these symptoms and other disorders (Prata *et al.*, 2009) the findings presented here may be secondary in nature to the neurocognitive and behavioural difficulties often described in preterm populations.

### Conclusion

Our findings highlight the impact of very preterm birth on mental health, lending support to the notion of a neurodevelopmental origin of psychopathology. These results further suggest that very preterm birth is a risk factor across a number of symptom domains and may not be limited to standard diagnostic boundaries. Further studies should focus on the investigation of known antecedents of psychopathology in very preterm children, such as emotion regulation problems (Treyvaud *et al.*, 2012). These results further suggest that early preventative interventions should extend to target individuals born very preterm.

## Acknowledgements

We thank our study participants for their continuing help. We also thank the National Institute for Health Research (NIHR) Biomedical Research Centre at South London and Maudsley NHS Foundation Trust. The study was funded by the Medical Research Council, UK (ref. MR/K004867/1).

The authors have no financial relationships relevant to this article to disclose.

